# Synthetically engineered IgG1 antibody Fc fragments presenting influenza A virus receptor sialic acid inhibit viral haemagglutination activity, but enhance virus replication in cultured A549 cells

**DOI:** 10.1101/2022.10.06.511107

**Authors:** Christina Vrettou, Patricia Blundell, Eleanor R Gaunt, Richard J Pleass

## Abstract

Many clinically important viruses, including influenza A, SARS-CoV-1, adenoviruses, and DNA tumour viruses such as Kaposi’s sarcoma herpesvirus use multivalent binding to sialic acid (SA) to infect cells, or to modulate immune responses through interactions with sialylated attachment factors that facilitate virus infectivity and/or host survival. Molecular scaffolds rich in SA that bind virions with high avidity may therefore be useful as anti-infective medicines. We generated a panel of 12 of these molecules using fragment-crystallisable scaffolds in CHO-S cells that are rich in SA. The viral surface protein of influenza A virus (IAV), haemagglutinin, binds SA for cell entry, and so we tested the activity of these compounds against this virus. Two of the sialylated Fc-molecules reduced IAV haemagglutination activity by up to 64-fold. However, the same molecules enhanced virus infectivity of A549 cultured cells. To explain the increased viral titres, we postulated that sialylated Fcs may be anti-inflammatory. However, sialylated Fc multimers were instead pro-inflammatory; they induced chemokine/cytokine responses from differentiated human THP-1 derived macrophages, including raised IL-8 and MIP-1α/β, that mimicked responses driven by universal type I interferon. Steric targeting of SA to block virus entry may therefore have unexpected effects in target cells that currently preclude their use for medical intervention.

## INTRODUCTION

Sialic acid (SA) linked to glycoproteins and gangliosides at the cell surface are used by many viruses as attachment receptors for host cell entry (1). Attachment to SA is mediated through receptor-binding proteins that are exposed on the surface of virus particles (1–3). Some SA-binding viruses are also equipped with neuraminidase (NA) or a sialyl-*O-*acetylesterase, which are receptor-destroying enzymes that can promote virus release from infected cells and prevent their re-entry into a cell that is already infected (1, 2).

Influenza A virus (IAV) assembles hundreds of haemagglutinin (HA) trimers on its surface to recognize SA-galactose linkages on target tissue (4, 5). The monovalent interaction between HA and a typical sialylated ligand is weak (mM range), but multivalency enhanced interactions allow firmer viral adhesion to the target cell at low concentration (6, 7). Molecules that bind HA with high avidity may therefore be useful as anti-infective medicines (6–8). Consequently, many studies have explored multivalent scaffolds to present SA to HA with the aim of blocking the interaction between virus and host receptors (6, 7, 9–14).

The potential for a range of different scaffolds to chemically attach to SA and block virus binding has been explored. These include antibodies, DNA, fullerenes, graphene, polyacrylamide, quantum dots, magnets, silver and gold nano particles; biocompatibility and potential toxicological liabilities of all of these mean that none have progressed to approval for use in humans (15, 16). Controlling the spatial distribution, type, and number of ligand-bearing units in these oligomers can also be sub-optimal and/or ill-defined, leading to reduced binding or promiscuous binding to irrelevant receptors. The lack of target specificity can lead to faster *in vivo* clearance rates and may also explain reported toxicities for such inhibitors (17, 18).

These limitations have driven the search for smaller, rationally designed scaffolds conjugated with SA (i.e. ‘sialylated’) to act as receptor decoys, for example rigid self-assembled peptide-nucleic acid complexes (6, 7). These have been used to show that smaller bivalent displays of the sialyl-LacNAc ligand (50–68Å between each sugar) are more effective at binding a single HA trimer and inhibiting haemagglutination by virus than longer scaffolds with SA residues separated by distances >100Å (6, 7). Longer scaffolds may nonetheless allow for inter-HA bridging, which may be desirable, as this may hinder the movement and clustering of HA around its target receptor (6, 7, 19).

Naturally sialylated scaffolds, including IgG-Fc, have also been explored for their therapeutic potential (20). Antibody Fc-fragments derived from the proteolytic cleavage of human IgG have been administered safely to children in the treatment of idiopathic thrombocytopenia (ITP) (21). Consequently, glycoengineering approaches to further enhance the therapeutic utility of the Fc for ITP, or other diseases for which the sialylated Fc plays a major role are actively being pursued by academics and pharmaceutical companies (19).

Here we explore the Fc of IgG1 as an alternative scaffold to template SA and, investigate its interactions with IAV HA. Electron micrograph analyses have shown that the average distance between two adjacent HA trimers to be ~101.7±0.6Å, and therefore within reach of both the proximal and distal SA’s on an Fc fragment *(Fig. 1A)* (6, 7, 19). We previously engineered a small library of 36 sialylated Fc molecules with different combinations of SA moieties and investigated their impact on IAV and influenza B virus (IBV) binding to human cells (22, 23). Altered sialic content was achieved by amino acid mutations introduced at N-linked sites that are not conventionally sialylated *(Fig. 1A)* (22–24). A small number of these compounds blocked haemagglutination by UV attenuated IAV and IBV in the high nanomolar concentration range (22, 23).

**FIGURE 1.**
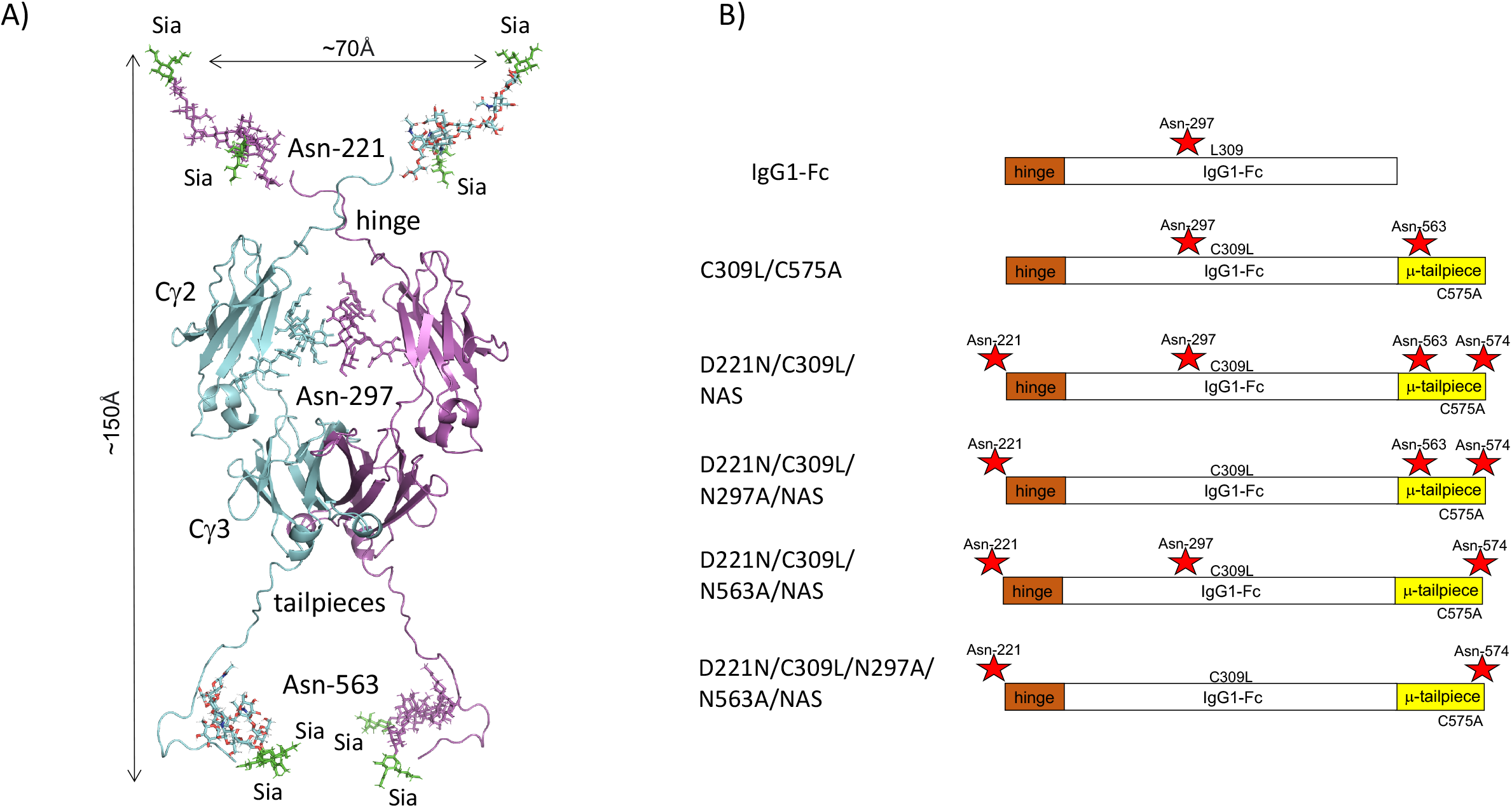
A model of sialylated Fc showing the position of critical N-linked glycans and generation of the NAS panel of mutants. (A) Structural model of the Fc with introduced Asn-221, Asn-297, and Asn-563 (D221N/C309L/C575A) containing at least eight SA (green) combining sites was modelled on the IgG1 structure (pdb 1HZH)(52). The N-linked glycan was attached using glycan entry 8388 in GlyProt (53) using torsion angles of 180, 340, 160, 280 and 90, 180, 160, 280 for the Asn-221 and Asn-563 attached sugars, respectively. The Asn-297 glycans are superimposed from the 1HZH IgG1 structure on which the dimer co-ordinates and final model was aligned using PyMol version 1.20 available at http://www.pymol.org/pymol. (B) Schematic showing the NAS panel of glycan mutants created by mutating the last three tailpiece amino acids from TCY to NAS. Note that these mutations remove Cys-575 and creates a NAS sequon for the attachment of N-linked carbohydrate. Red stars indicate the hinge Asn-221, the Cγ2 Asn-297, and the tailpiece Asn-563 and Asn-574 glycan sites.

Binding of sialylated Fc fragments to HA is driven by the hinge-attached sialylated glycan at Asn-221 (*Fig. 1A*) (22, 23). Conversely, a tailpiece Asn-563 glycan that is rich in SA is not required (24, 25). Modelling the interaction of the sialylated Fc with HA suggested that the orientation of the Asn-563 glycan may not be optimal for interactions with HA (*Fig. 1A*) (19).

Here we describe the creation of a further set of Fc-mutants (the NAS series) in which an additional N-linked glycosylation sequon (N-X-T/S hence NAS) is inserted at Asn-574 (*Fig. 1B*), a location on the tailpiece predicted from our modelling data to be more accessible to HA (19). While binding to live infectious IAV was improved through the addition of Asn-574 (as demonstrated by haemagglutination inhibition assays), we observed that two NAS mutants significantly enhanced the ability of IAV to infect A549 lung epithelial cells. Cytokine and chemokine array profiling of macrophages stimulated with the NAS mutant that most potently inhibited IAV-induced haemagglutination highlighted overlapping features with the type I interferon response. The results highlight potential drawbacks and possible solutions to using sialylated Fc scaffolds in the treatment of viruses while opening up new areas for study and potential applications.

## MATERIALS AND METHODS

### Generation of Fc fragments

To insert an extra N-linked glycosylation sequon (N-X-S/T) at a more distal position on the tailpiece to previously engineered mutants (22–24) (Fig. 1A), while maintaining the lack of potential for covalent bonding through Cys-575, we modified the last three tailpiece amino acids (Eu amino acid numbers 574, 575, and 576) from TCY to NAS (*Fig. 1B).* The NAS series of mutants (*Fig. 1B*) were made by PCR using primers 5’-CGGACTGGGACGAACGAGTTGAGA-3’ and 3’-GGCCAGCTAGCTCAGGAGGCGTTGC-5’ with the previously described set of glycan-adapted CL309/310LH plasmids as template and under previously described conditions (22, 23). To verify incorporation of the desired mutation and to check for PCR-induced errors, the open reading frames of the new mutants were sequenced on both strands using previously described flanking primers (22, 23). CHO-K1 or HEK-293F cells (European Collection of Authenticated Cell Cultures) were transfected with plasmids using FuGene (Promega), and Fc-secreting cells cloned, expanded, and the proteins purified as previously described (22, 23). To generate the quantities of Fc fragments required for virus neutralisation studies, manufacture of selected Fc mutants was outsourced to Abzena Ltd for synthesis in their suspension-adapted CHO-S cells (originally CHO-K1 received from ATCC and adapted to serum-free growth in suspension culture). Constructs were gene-synthesized and cloned into the proprietary expression vector, pEVI-5, using conventional (non-PCR based) methods. Plasmid DNA was prepared, cells were transfected with eviFect, a proprietary transfection reagent, and cells were grown after transfection in eviMake, an animal component-free, serum-free medium. Supernatants were harvested by low-speed centrifugation and subsequent filtration (0·2-μm filter). Fc fragments were purified from cell culture supernatant using a 5 ml HiTrap Protein G HP column (GE Healthcare). The supernatants for each mutant were processed as two separate batches. Each time, the column was washed using 20 mM sodium phosphate pH 7·2 containing 150 mM NaCl and protein was eluted using 0·1 M glycine (pH 2·7). Collected fractions were neutralized using 1 M Tris pH 9·0 and then buffer-exchanged using Zeba™ Spin 7 K MWCO, 10 ml Desalting Columns (Thermo Fisher, Loughborough, UK), into 10 mM sodium acetate pH 5·6 buffer containing 100 mM NaCl. The final samples were then filter-sterilized before quantification at A280 nm using an extinction coefficient Ec (0·1%) based on the predicted amino acid sequence. Purified proteins were then analysed by SDS-PAGE (*Fig. 2E,F*) and analytical size-exclusion HPLC (*Fig. 2G,H*). All the glycan-adapted Fcs, with the exception of the C309L/N297A/N563A/NAS mutant, could be made in-house or by Abzena to the g/L yields required for use in cell-based assays.

**FIGURE 2.**
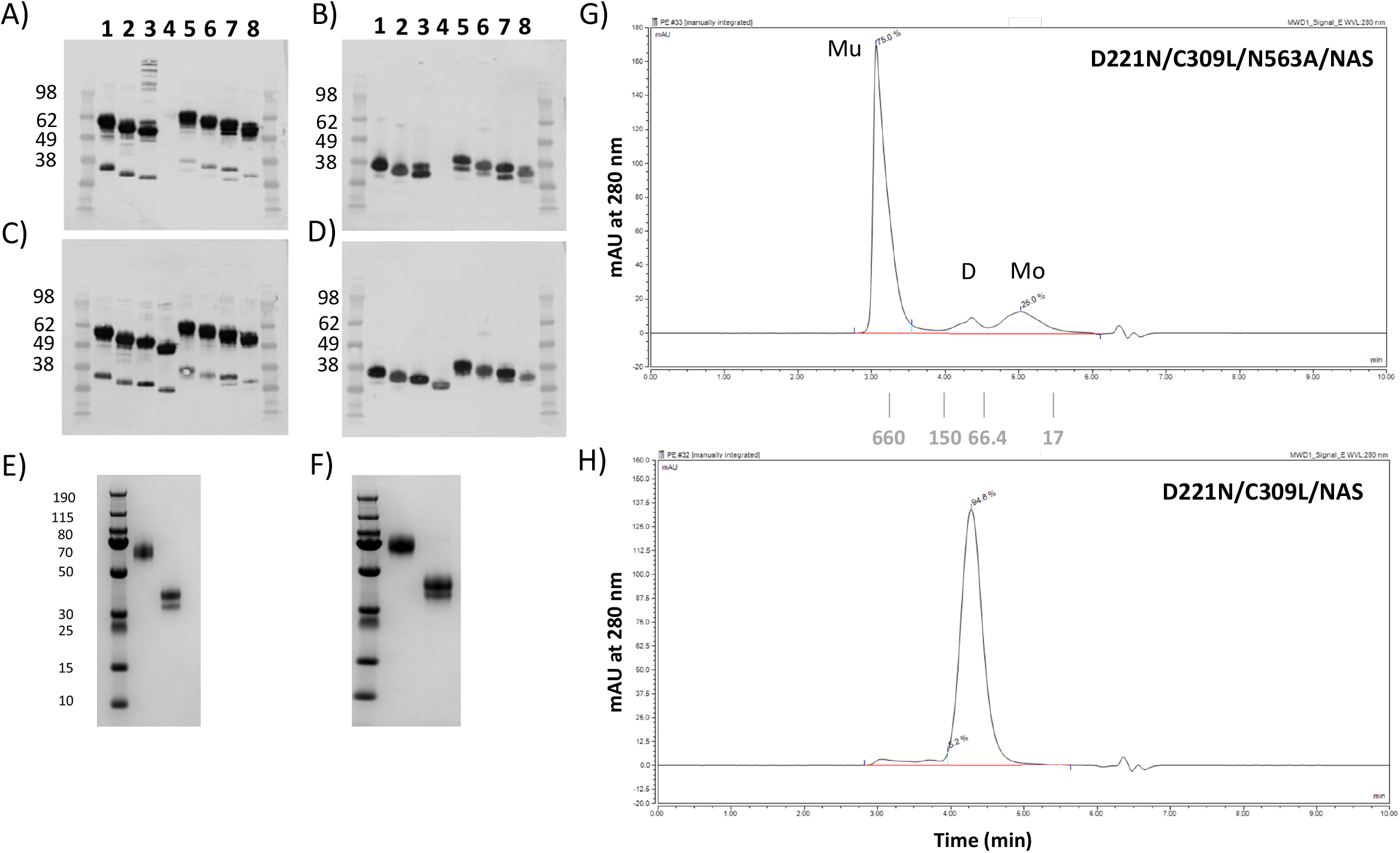
Characterisation of the Fc NAS mutants by SDS-PAGE and size-exclusion chromatography. (A) Non-reducing and (B) reducing conditions for 2 μg of Fc purified from CHO-K1 cell culture supernatant. (C) Non-reducing and (D) reducing conditions for 2 μg of Fc purified from HEK-293F cell culture supernatant. Lane 1 = C309L/NAS; lane 2 = C309L/N297A/NAS; lane 3 = C309L/N563A/NAS; lane 4 = C309L/N297A/N563A/NAS which was not expressed; lane 5 = D221N/C309L/NAS; lane 6 = D221N/C309L/N297A/NAS; lane 7 = D221N/C309L/N563A/NAS; lane 8 = D221N/C309L/N297A/N563A/NAS. Proteins were run into 4-8% gradient acrylamide gels, transferred to nitrocellulose, and blotted with anti-human IgG Fc (Jackson labs). (E) Coomassie stained SDS-PAGE gel of 5 μg of D221N/C309L/N563A/NAS protein expressed by Abzena in CHO-S cells run under non-reducing and reducing conditions respectively. (F) Coomassie stained SDS-PAGE gel of 5 μg of D221N/C309L/NAS protein expressed by Abzena in CHO-S cells run under non-reducing and reducing conditions respectively. SEC-HPLC chromatograms as described in methods for the Abzena purified Fc mutants (G) D221N/C309L/N563A/NAS and (H) D221N/C309L/NAS respectively. Molecular weights are shown in kDa.

### Cells and viruses

MDCK-SIAT (26), human embryonic kidney (HEK)-293T and A549 cells (Sigma, Glasgow, UK) were grown in Dulbecco’s Modified Eagle Medium (DMEM) (Sigma) supplemented with 10% v/v foetal calf serum (FCS) (Thermo Fisher Scientific, Waltham, MA) and 1% v/v penicillin/ streptomycin (Thermo Fisher Scientific). Cells were maintained by twice-weekly passage and checked for mycoplasma contamination regularly using a MycoAlert mycoplasma detection kit (Lonza, UK). IAV strain A/California/04/2009 (H1N1) (‘Cal04’) was rescued using reverse genetics as described elsewhere (27, 28). Briefly, 70% confluent HEK293T cells in 6 well plates were transfected in Opti-MEM reduced serum medium (Thermo Fisher) with 250 ng of 8 plasmids, each corresponding with a segment of viral genome, and 4 μl Lipofectamine 2000 transfection reagent (Thermo Fisher) per well. From here on cells were grown at 35°C. After 24 hr, medium was replaced with serum free medium containing 1 μg/ml tosyl phenylalanyl chloromethyl ketone (TPCK)-treated trypsin (Sigma) and 0.14% BSA (w/v) (Sigma). At 72 hr post-transfection, supernatants were collected and 100 μl was inoculated into the allantoic fluid of an embryonated hen’s egg. Dekalb White fertilised eggs, purchased from Henry Stewart (Norfolk, UK), were incubated at 37.5°C, 50% relative humidity until infection at day 11. 100 μl virus was injected into the allantoic fluid and holes were sealed with sticky tape under sterile conditions. For preparation of virus stocks, infected eggs were left for 72 hr at 35°C prior to allantoic fluid harvest, which was spun down for 5 min at 800*g* to pellet cellular debris, and supernatant was aliquoted and used as neat virus stock. Virus titres were determined by plaque assay.

### Plaque Assays

MDCK-SIAT cells were seeded in 12 well plates. Upon achieving confluency, cells were washed twice with serum free media. Ten-fold serial dilutions of virus stock or infection supernatant were made using serum free media. 450 μl of each dilution (10^-1^ to 10^-6^) was added and incubated for 1 hr. 1 ml overlay comprising of 34% v/v of 2.4% w/v cellulose (Sigma), 66% v/v serum free media, 2% v/v of 7% w/v BSA (Sigma) and 1 μg/ml tosyl phenylalanyl chloromethyl ketone (TPCK)-treated trypsin (Sigma) was then added on each well. Plates were returned to the 35°C and 5% CO_2_ and left for 72 hr. Next, 1 ml of neutral buffered formalin (NBF) was added / well and left for at least 30 minutes. NBF was then replaced with 0.1% toluidine blue Sigma) for at least 30 min, after which time plaques were counted.

### Haemagglutination assays

1% v/v red blood cells (erythrocytes) were prepared by washing whole chicken or human blood, thrice, with PBS and spinning at 1500 rpm for 5 min, with a last step of centrifuging at 2500 rpm for 10 min, before re-suspending at 1% v/v in PBS. For the HA assays, 50 μl of PBS was added to appropriate wells of a 96-well plate. Next, 50 μl of 4 μM test compound was added on the first well to give a 1 in 2 dilution. Two-fold serial dilutions were made by transferring 50 μl across the plate, discarding the final 50 μl. 50 μl of 4HA units of Cal04 virus was then added per well and left to incubate on ice for 30 min. Cal04 in the absence of any compound was included as control. Thirty minutes after the addition of virus, 50 μl of erythrocytes was added and left for 30 min at room temperature, before recording results. The determination of the 4HA units was done by previously titrating the virus on a separate HA assay in which 4HA units was considered to be the dilution at two wells ahead of the last well where complete haemagglutination occurs (i.e, 4-fold the minimum required amount of virus for haemagglutination to be achieved).

### Virus infections

A549 cells at ~90% confluency in 24 well plates were washed twice and complete media was replaced with serum free media. Cells were infected with Cal04 virus at a multiplicity of infection (MOI) of 3 and each of the testing compounds was used at final concentration of 2 μM. The virus was mixed with each compound in a total volume of 200 μl serum free medium, and left to incubate on ice for 30 min. Virus-compound mixes were then added to cells in technical duplicate, incubated for 1 hr, and then cells were washed and overlaid with serum free media for 24 hr. Cells incubated with only virus or only compound were used as controls. The supernatants were then harvested and used in plaque assays for quantification of the production of infectious virus after a single infection cycle.

### Cytokine profiling

THP-1 monocytes (human acute monocytic leukemia cell line), differentiated into macrophages using PMA, were used to assess cytokine induction by compounds. THP-1 cells were gifted by the lab of Dr Musa Hassan, Roslin Institute. Cultures were maintained in RPMI1640 (Sigma) media enriched with 10% FBS, 1X Glutamax and 1% Pen/Strep and maintained at densities between 1×10^5^/ml and 1×10^6^/ml. For their differentiation into macrophages, 200 μl/well of cells were plated into 96 well plates, at 1×10^5^ per well with 100 nM PMA. Cells were left to differentiate for 72 hr at 37°C and 5% CO_2_, washed twice with complete media and left without PMA for 24 hr before assaying. For the cytokine assays, cells were treated for 1 hr with either 10^4^ units universal type I interferon (Stratech, 11200-1-PBL), test compound (2 μM) or a mixture of both interferon and test compound at the same concentrations as used individually. Cell supernatants were then harvested and assayed for cytokines by Human Cytokine Array (Proteome Profiler, R&D systems, ARY005B), according to the manufacturer’s instructions.

### Statistical analyses

Figures were made and ratio-paired t tests were performed in GraphPad Prism software, with a *p* value <0.05 considered to be significant.

## RESULTS

### Moving the IgM tailpiece N-linked glycosylation site from Asn-563 to Asn-574 does not disrupt multimerization of the IgG1-Fc

After expression in CHO-K1, CHO-S or HEK-293F cells and protein G affinity purification, the integrity and purity of each of the NAS Fc mutants was confirmed by SDS-PAGE. All the mutants could readily be made with the exception of the C309L/N297A/N563A/NAS mutant in CHO-K1 cells (*Fig. 2, A,B).* All the mutant Fcs migrated on SDS-PAGE with the expected molecular weight for their glycosylation or disulfide bonding states (*Fig. 2).*

Earlier studies in CHO-K1, CHO-S and HEK-293F cells demonstrated that all Fcs in which the tailpiece Asn-563 glycan was substituted for alanine ran as non-covalent multimers in solution when examined by size-exclusion HPLC (22–24). Removal of the Asn-563 and insertion of a new glycosylation site further down the tailpiece at Asn-574 (as in D221N/C309L/N563A/NAS) resulted in the formation of 75% multimers with molecular weights of ~660 kDa approximating Fc dodecamers (*Fig. 2G).* The removal of the bulky Asn-563 glycan exposes hydrophobic amino acids in the Fc tailpiece that facilitate multimerization despite the presence of the new glycan inserted at Asn-574. Maintaining both the tailpiece Asn-563 and Asn-574 glycans (as in the D221N/C309L/NAS mutant) leads to molecules that run as ~5.2% multimers (~660 kDa), ~94.8% dimers (~110 kDa), and few to no observable monomers (*Fig. 2H).* The data show that the number and positioning of the attached N-linked glycan on the tailpiece contributes to the type of multimer formed.

### The NAS series of human IgG1 Fc mutants inhibit haemagglutination by influenza A virus

To test if the insertion of a distally located glycan at position Asn-574 leads to enhanced inhibition of SA binding by IAV, we screened the new NAS panel of mutants using the World Health Organization haemagglutination inhibition assay (HIA) (29) to quantify inhibitory titres for each mutant against an infectious Cal04 H1N1 influenza human-derived isolate (*Fig.3*). The NAS mutants in general were superior to their matched controls (without Asn-574) at inhibiting virus agglutination of human (*Fig. 3A*) or chicken (*Fig. 3B*) erythrocytes. For example, as little as 0.06 μM of the D221N/C309L/N563A/NAS mutant inhibited haemagglutination by Cal04 whereas 2 μM of the matched control D221N/C309L/N563A/C575A was without effect, irrespective of the cell line used to manufacture the Fc or the source of erythrocytes used in the HIA. Mutants that had previously shown efficacy in HIA using UV irradiated New Caledonia/20/99 H1N1 virus, e.g., D221N/C309L/N297A/C575A and D221N/C575A (22, 23) were capable of inhibiting agglutination by infectious Cal04 virus using chicken erythrocytes (*supplementary Fig. 1*) but not human erythrocytes (*Fig. 3A*), a finding we attribute to the known differences in sialylation between chicken and human erythrocytes (30). Only one of the parental set of Fcs (D221N/C309L/N563A/C575A) showed any activity at all with human erythrocytes, a finding that may be explained by the use of live virus rather than UV-inactivated viruses published previously (22, 23).

**FIGURE 3.**
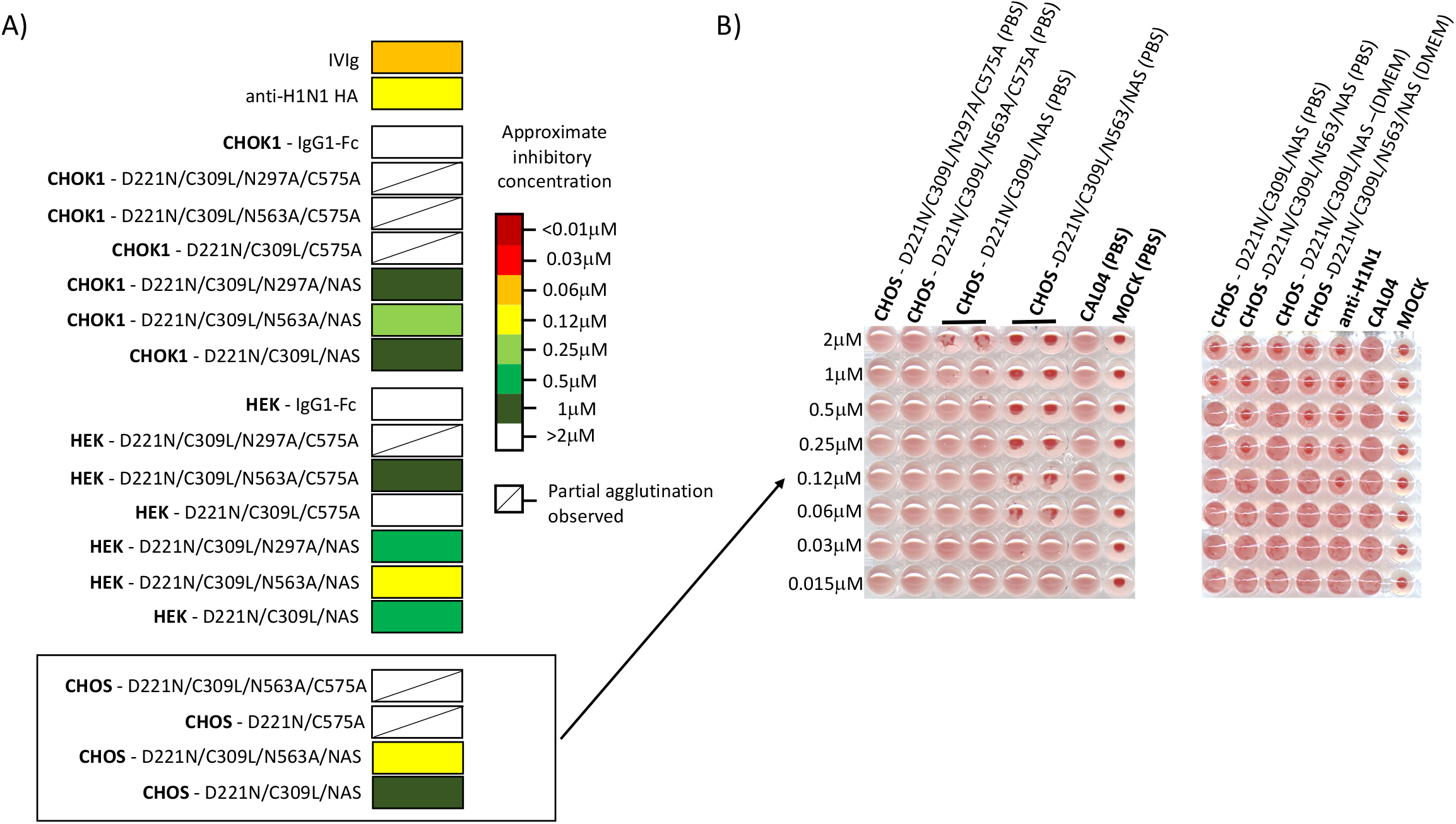
Fc NAS glycosylation strongly enhances haemagglutination inhibition. 4 HA units of influenza A Cal04 virus was incubated with titrated amounts of Fc glycan mutants and added to (A) human O+ erythrocytes and allowed to sediment on ice for 1 h. Result are representative of two independent experiments which for simplification are presented graphically. (B) Wet experiments with chicken erythrocytes in either PBS or DMEM are shown for selected mutants made by Abzena (boxed). Non-agglutinated erythrocytes form a small halo. Two independent experiments are shown.

### NAS mutants that inhibit agglutination of erythrocytes by IAV enhance infectivity of A549 cells irrespective of valency

We selected two of the strongest haemagglutination-inhibiting NAS mutants (D221N/C309L/NAS and D221N/C309L/N563A/NAS) for manufacture to scale in CHO-S cells and tested their ability to inhibit virus infection in the human lung carcinoma A549 cells. The A549 cell line is commonly used to study influenza virus biology (31). Contrary to expectation, 2 μM of either the dimeric D221N/C309L/NAS or multimeric D221N/C309L/N563A/NAS Fc-mutant significantly enhanced production of infectious virus in A549 cells by 30-fold and 17-fold respectively (*Fig. 4A*).

**FIGURE 4.**
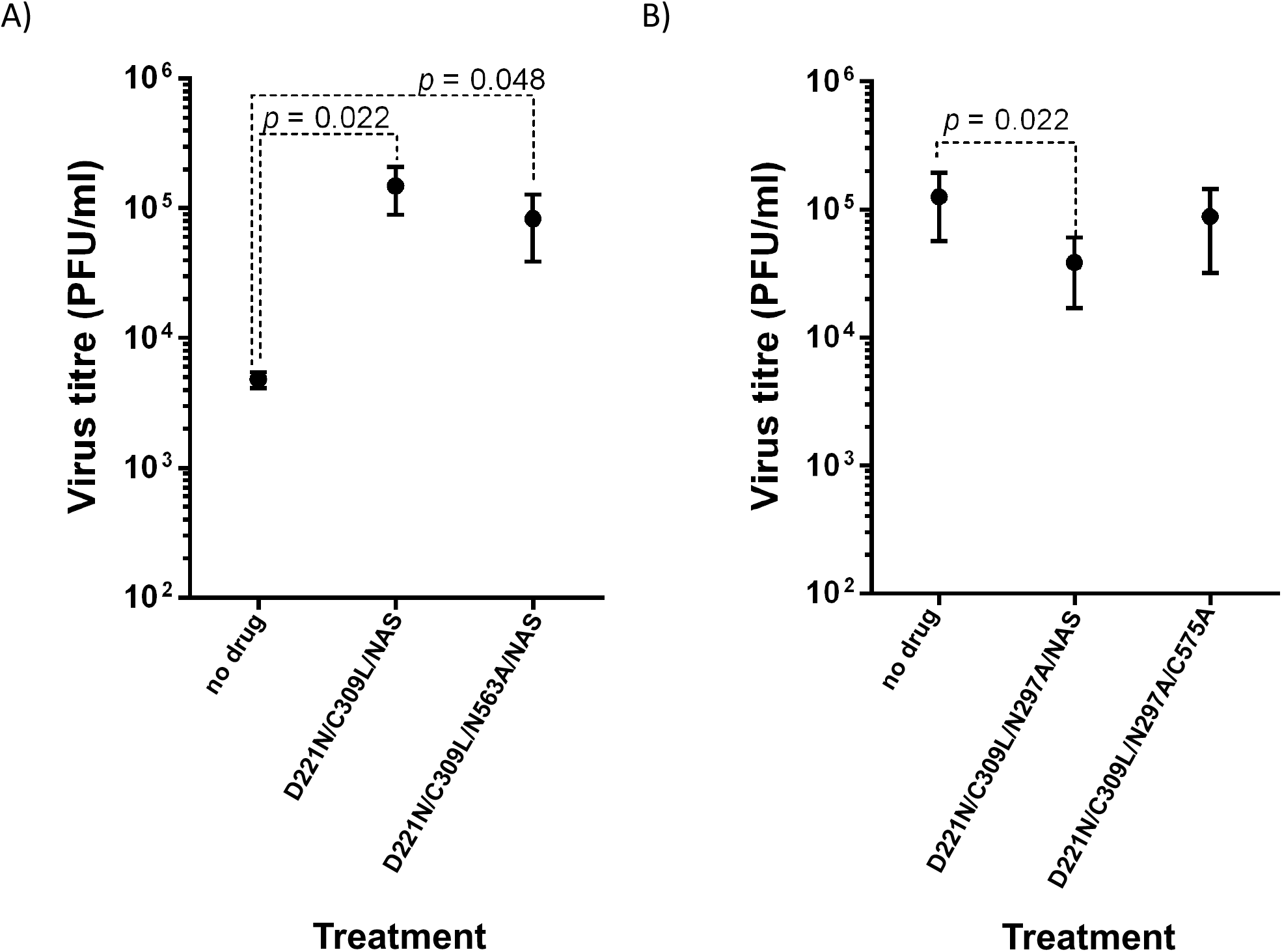
Fc mutants that inhibit haemagglutination strongly enhance A549 cell infection with IAV in an Asn-297-dependent manner. 2 μM of each Fc NAS mutant was incubated with 3 x 10^5^ PFU Cal04 virus and left to bind on ice for 30 mins prior to adding to A549 cells at a MOI 3.0. After 1 h plates were washed, and fresh media added and incubated for a further 24 hrs. Supernatants were titrated by plaque assay. (A) The impact of D221N/C309L/NAS and D221N/C309L/N563A/NAS mutants on Cal04 IAV infectivity of A549 cells. (B) The impact of paired Asn-297 deficient mutants D221N/C309L/N297A/NAS and D221N/C309L/N297A/C575A on Cal04 IAV infection in A549 cells did not alter virus production. Bars show standard error of the means.

### Asn-297 deletion leads to NAS mutants with reduced infectivity of A549 cells

We hypothesised that the enhanced infectivity of A549 cells seen as a result of treatment with the D221N/C309L/NAS and D221N/C309L/N563A/NAS Fc-mutants may arise from cross-bridging of bound virus to endocytic receptors found on A549 cells. Previous studies have shown that removal of the conserved glycan at Asn-297 abolishes binding of these Fc-fragments to classical FcγRs, C1q and lectin receptors (22), and inhibits their functionality (32). To test this hypothesis, we generated two further N297A mutants in CHO-S cells, including D221N/C309L/N297A/NAS and its matched control D221N/C309L/N297A/C575A; the former mutation being within the parent D221N/C309L/NAS molecule that enhanced infectivity (*Fig. 4A*). The D221N/C309L/N297A/NAS mutant this time led to significantly reduced production of infectious virus, and the D221N/C309L/N297A/C575A mutant did not alter virus production, consistent with our hypothesis (*Fig. 4B*).

### The NAS mutant D221N/C309L/N563A/NAS drives IL-8 and MIP-1α/β production from differentiated THP-1 cells

Although Fc compounds that facilitate virus entry may not make ideal antivirals, they may be practically useful to enhance virus uptake by cells during virology assays, and also may be therapeutically useful as immune modulators of virus-mediated disease (19). To examine the impact of Fc compounds on antiviral signalling (relevant for both applications), we incubated THP-1 derived human macrophages with either D221N/C309L/N563A/NAS and/or IFN-α (as a positive control) and measured the relative production of 36 human cytokines released into culture supernatant over a 1 h period (*Fig. 5*). Seven of the 36 arrayed cytokines/chemokines were induced after stimulation of THP-1 cells with either IFN-α, of which 6 were also induced by the D221N/C309L/N563A/NAS mutant (*Fig. 5*). A marked induction of MIP-1α/β and IL-8 resulted from treatment of cells with D221N/C309L/N563A/NAS. MIP-1α/β and IL-8 are both chemotactic cytokines secreted by macrophages to recruit leukocytes and neutrophils respectively. In contrast, IFN-α treatment alone led to raised IL-1ra and MIF of which only IL-1ra levels could be reduced by co-culture with D221N/C309L/N563A/NAS (*Fig. 5*). Other proinflammatory cytokines, including IL-1α/β, IL-6, IL-12, IL-18, IFN-γ, tumour necrosis factor (TNF), and granulocyte-macrophage colony stimulating factor (GM-CSF) were notable by their absence (*Fig. 5*). Overall, mutant D221N/C309L/N563A/NAS induced cytokine expression in a profile similar, but not identical to, type I interferon in macrophages.

**FIGURE 5.**
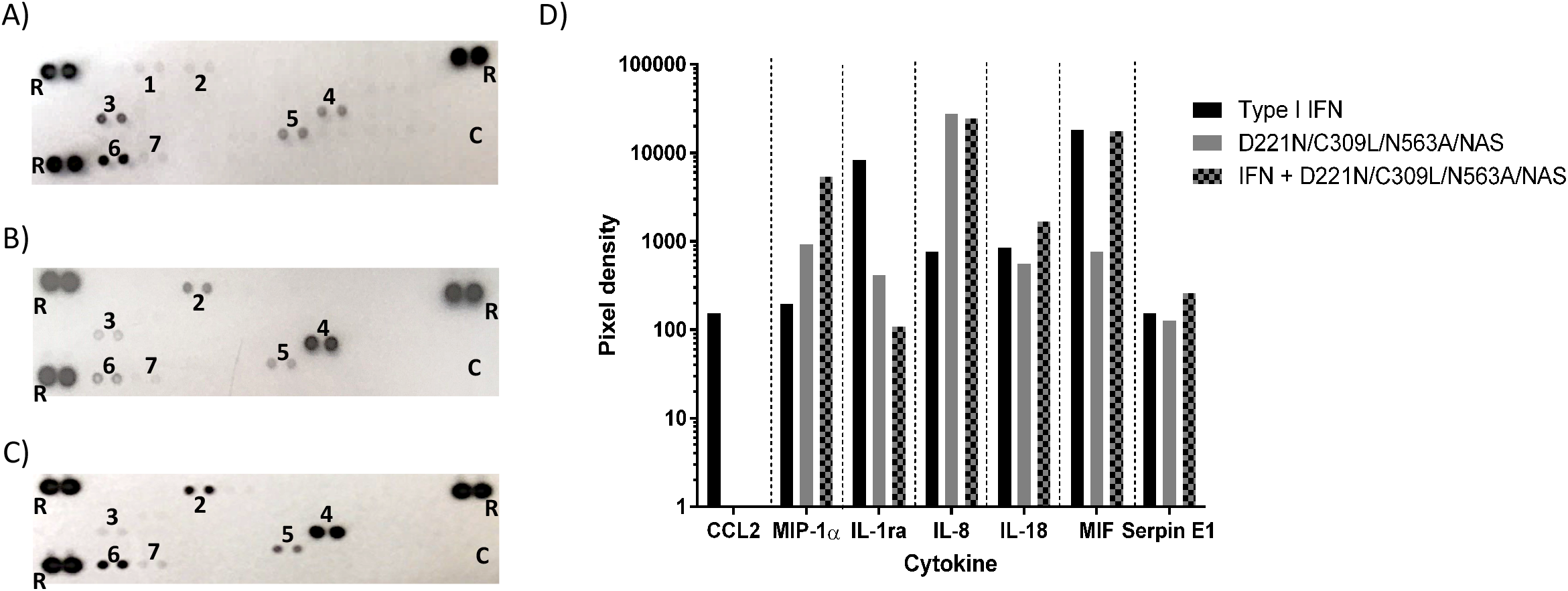
Fc mutant induction of cytokines in macrophages is highly concordant with cytokine induction by type I interferon. Wells were treated for 1h with: A) type I interferon, B) D221N/C309L/N563A/NAS-Fc, or C) type I interferon and D221N/C309L/N563A/NAS-Fc. Pairs of spots represent analyte captured from culture supernatants and detected by antibodies on the membrane and are numbered as follows, 1 = CCL2/MCP-1; 2 = MIP-1α/MIP-1β; 3 = IL-1ra/IL-1F3; 4 = IL-8; 5 = IL-18/IL-1F4; 6 = MIF; 7 = Serpin E1/PAI-1; R = Reference spots, C = Negative control, ND = not detected. D) Mean spot pixel densities determined using a flatbed scanner and image J software.

## DISCUSSION

We previously published that Fc-fragments with SA attached at Asn-221 and Asn-563 can inhibit agglutination of human erythrocytes by two genetically divergent influenza A (H1N1) and influenza B (Hong-Kong) inactivated viruses using the WHO standard HIA (22, 23). Although HIA is a good correlate for the potential *in vivo* efficacy of these reagents, the findings need to be supported with more demanding virus neutralisation experiments with permissive cell lines that better mimic the human lung infected with live virus.

Here we generated a new panel of glycosylated Fc-mutants with improved HA binding by moving the Asn-563 located glycan to a more exposed position (amino acid 574) on the Fc (*Fig. 1B* & *Fig. 3*). The sugars attached at Asn-563 prevent non-covalent interactions of underlying hydrophobic amino acids e.g., Val-564 & Val-567, that facilitate Fc clustering (24). The replacement of Asn-563 with Asn-574 likely enhances haemagglutination inhibition by the D221/C309L/N563A/NAS mutant through high avidity interactions with virus as a consequence of observed multimer formation (*Fig. 2G*), as dimers that retain both the Asn-563 and Asn-574 glycans bound virus weakly (*Fig. 2H* & *Fig. 3B*).

Despite inhibiting haemagglutination by Cal04 IAV, the D221/C309L/N563A/NAS and D221/C309L/NAS Fc-mutants markedly enhanced virus propagation of Fc-opsonised IAV in A549 cells (*Fig. 4A*). Interestingly, the D221N/C309L/NAS mutant that did not bind virus strongly in HA assays (Fig. 3B) and existed mostly in dimeric form was particularly effective at facilitating virus entry and replication in A549 cells (*Fig. 4A*).

One hypothesis to explain our observation is that sialylated Fc-fragments can cross-bridge IAV virions to underlying endocytic receptors found on or in A549 cells that facilitate virus entry (*Fig. 6*). Our previous studies have shown that removing glycosylation with the N297A mutation abolishes interactions of D221N/C309L/N297A/C575A with FcγRs and significantly reduces binding to multiple carbohydrate binding receptors including, Siglec-1, Siglec-2, Siglec-3, CD23, dectin-1, dectin-2, DC-SIGN, CLEC-4D, DCIR, MBL and MMR (22, 23). We therefore tested two further mutants, D221N/C309L/N297A/NAS and its matched control D221N/C309L/N297A/C575A, in which the conserved Asn-297 glycan was removed by mutation to alanine. Both of these mutants did not enhance viral replication, suggesting a role for classical FcγRs and/or lectin receptors in enhancing the infectivity of A549 cells (*Fig. 4B*). However, A549 cells do not express classical FcγRs (33). Other possible endocytic receptors for Fc-opsonised virus include the neonatal Fc-receptor (FcRn) and the universal tripartite motif containing-21 (TRIM21). However, interactions of both FcRn (34) and TRIM21 (35) for the Fc of IgG are unaffected by removal of the N-glycan structure attached to Asn-297 in the Fc. The epidermal growth factor receptor (EGFR) is activated by IAV and transmits cell entry signals into A549 cells (36). As EGFR is not known to bind SA or the Fc, our current hypothesis is that IAVs coated in sialylated-Fc cross-bridge to unidentified carbohydrate receptors on A549 cells.

**FIGURE 6.**
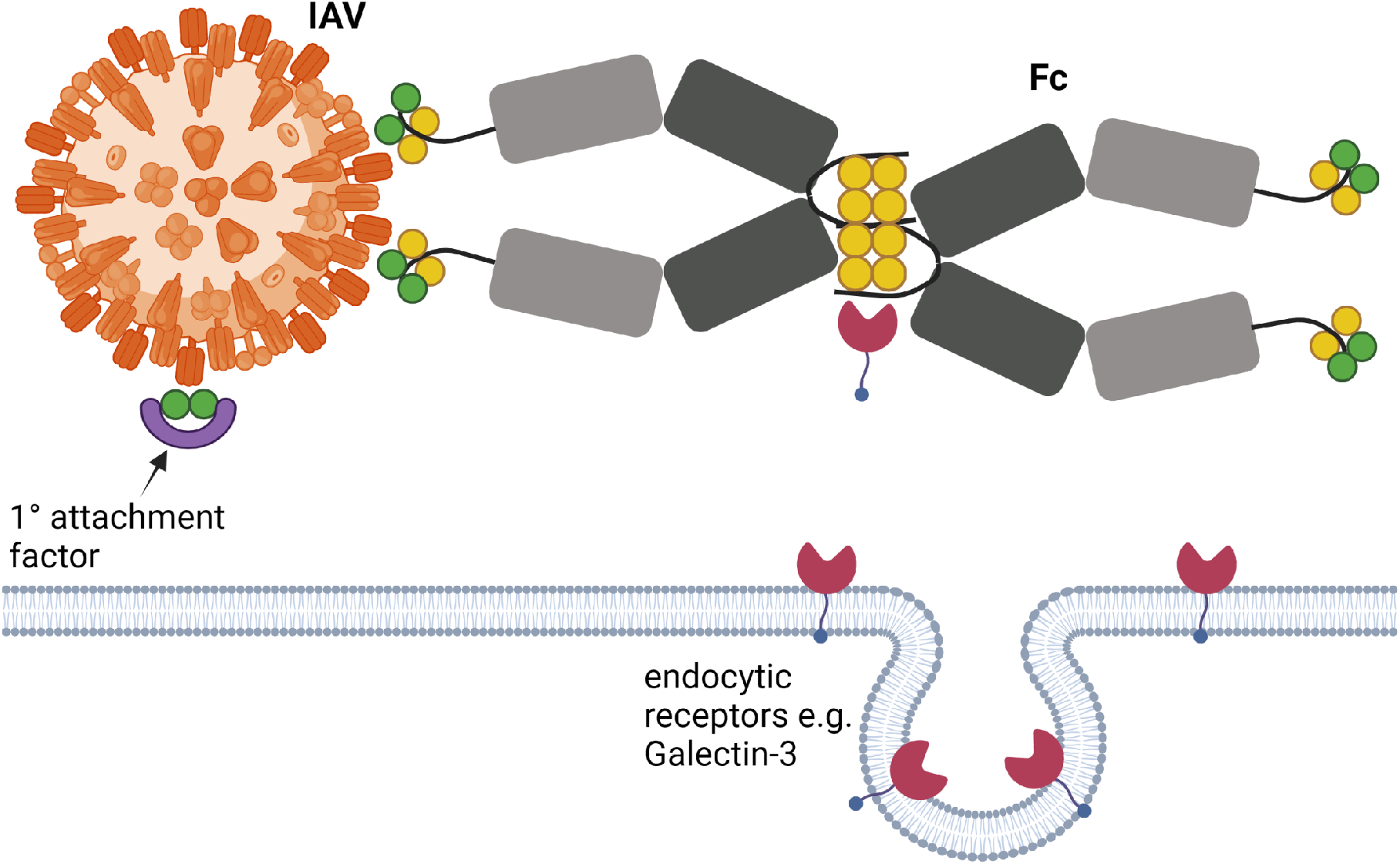
Model of the proposed cross-bridging of influenza virus to endocytic receptors in A549 cells by the Fc. The sialylated Fc (grey) is bound by haemagglutinin (HA) expressed on the surface of influenza A virus particles (orange). Because we have shown that Asn-221 and Asn-563 containing Fc is rich in galactose (22, 23, 32), we hypothesise that Fc bound to virus through SA (dark green circles) can cross-bridge galactose (yellow circles) to endocytic receptors, including galectin-3 on A549 cells. Galectin-3 (red triangles) is abundantly expressed by A549 cells (37, 38), is phagocytic (44), and is already known to bind galactose residues found on the Fc of human IgG1 (39).

We searched published A549 proteomes to find likely candidate receptors known to bind the IgG-Fc (37, 38). The only abundant protein in A549 cells potentially capable of binding D221N/C309L/NAS and D221/C309L/N563A/NAS that we identified was Galectin-3, a soluble protein that regulates inflammatory responses and binds IgG1 immune-complexes rich in galactose (39). Galectin-3 is also known to regulate IAV infection (40). Although we know that glycans attached at Asn-221 and Asn-563 are richly decorated in galactose (22, 23), more work is required to determine if Galectin-3 is involved in enhancing the infectivity of viruses opsonised with either D221N/C309L/NAS or D221N/C309L/N563A/NAS.

We observed that Fc-mutants (e.g., D221N/C309L/N563A/NAS) that lack glycosylation at Asn-563 have a propensity to multimerise (*Fig. 2G*), a physical property that although facilitating stronger binding to HA (*Fig. 3*) may also enable higher avidity interactions with activating FcγRs (41). Cross-linking of FcγRs and/or Galectin-3 on monocytes/macrophages allows for the production of pro-inflammatory cytokines including IL-1β, TNFa, IL-6 and IL-8, that can result in immunogenicity-related adverse events (42–45). For instance, a Phase I trial of the anti-CD28 mAb TGN1412 resulted in immune-mediated cytokine storms, highlighting the importance of assessing the likelihood of such events arising with glycan-modified Fc-fragments (46). Here we show that D221N/C309L/N563A/NAS drives an atypical pro-inflammatory IFN-α-like response from differentiated human THP-1 cells, marked by elevated IL-8 and MIP-1α/β production and undetectable levels of other pro-inflammatory cytokines like IL-1, IL-6, IL-12, IL-17, IL-18, TNF-α and IFN-γ (*Fig. 5*). As IFN -α therapy is used in the treatment of certain viruses and cancers (47, 48), including glioblastomas (49), these findings may partly explain our earlier observations on the efficacy of N563A Fc-mutants at controlling immune-mediated demyelination by glial cells (32).

These data provide a basis for the continued preclinical evaluation of the variably glycosylated Fc in order to support their advancement to clinical studies of viral and/or neurological diseases. For example, further modifications to the N297A mutants may generate molecules that potently inhibit virus infection yet are not overtly pro-inflammatory. While it was unexpected that the ability of the Fc-NAS mutants to block viral binding of erythrocytes would translate to increased viral uptake in cultured cells, this may be useful to enhance virus infection assays. For example, the Cal04 strain does not infect A549 cells well; high MOI infections only result in infection of a low proportion of cells in a monolayer, as evidenced by confocal microscopy (50) and flow cytometry (51). Improving the proportion of cells infected may facilitate host-pathogen interaction studies and, overcome the limitations that are currently prohibitive of large-scale studies such as genome-wide CRISPR knockout screens.

## ACKNOWLEDGEMENTS

All authors were supported by a Bloomsbury SET Award. EG is supported by a BBSRC Institute Strategic Programme grant (BB/P013740/1) and a Wellcome Trust/ Royal Society Sir Henry Dale Fellowship (211222_Z_18_Z). The funders had no role in study design, data collection and analysis, decision to publish, or preparation of the manuscript. For the purpose of open access, the author has applied a CC BY public copyright licence to any Author Accepted Manuscript version arising from this submission. We are grateful to the labs of Dr Edward Hutchinson (MRC-University of Glasgow Centre for Virus Research, UK), and Professor Paul Digard and Dr Musa Hassan (The Roslin Institute, Edinburgh University, UK) for the provisions of plasmids and cells. We are grateful to Dr Colin Sharp, the Roslin Institute, for preparation of virus stocks. We are indebted to the staff of the Roslin Institute Central Services, and Facilities, Financial and Legal teams. We thank Abzena PLC for manufacturing the molecules described and for providing the data shown in Fig. 2E-H.

## CONFLICT OF INTEREST

Abzena provided no financial support. R.J.P. and P.A.B. declare that the molecules discussed within are subject to ongoing patent applications. The other authors have no financial conflicts of interest.

## AUTHOR CONTRIBUTIONS

P.B. generated the NAS series of mutants, C.V. did the haemagglutination, plaque and cytokine assays, R.J.P. and E.R.G. wrote the manuscript. All the authors read drafts and contributed to the final version.

## SUPPORTING INFORMATION

Additional supporting information may be found online in the Supporting Information section.

## Notes

**Funding information**, Bloomsbury SET Impact Connector Programme – Ref: 12RP-LSTM

### Competing Interest Statement

The authors have declared no competing interest.

