## Supplemental Fig 1 for "Synthetically engineered IgG1 antibody Fc fragments presenting influenza A virus receptor sialic acid inhibit viral haemagglutination activity, but enhance virus replication in cultured A549 cells"

### Slide 1
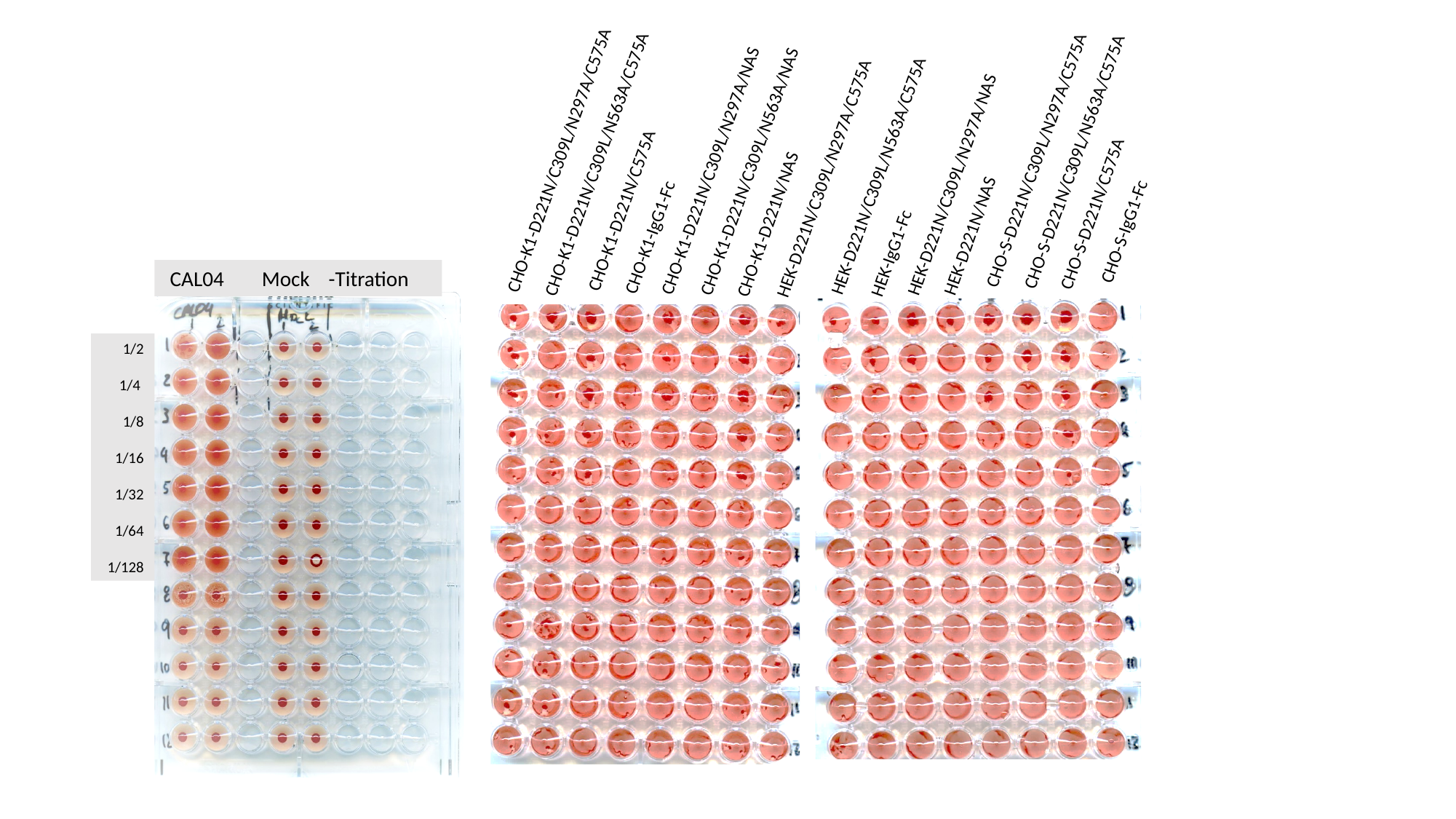

CHO-K1-D221N/C309L/N297A/C575A
CHO-S-D221N/C309L/N297A/C575A
CHO-S-D221N/C309L/N563A/C575A
CHO-K1-D221N/C309L/N563A/C575A
CHO-K1-D221N/C309L/N297A/NAS
CHO-K1-D221N/C309L/N563A/NAS
HEK-D221N/C309L/N563A/C575A
HEK-D221N/C309L/N297A/C575A
HEK-D221N/C309L/N297A/NAS
CHO-K1-D221N/C575A
CHO-S-D221N/C575A
CHO-K1-D221N/NAS
CHO-S-IgG1-Fc
HEK-D221N/NAS
CHO-K1-IgG1-Fc
HEK-IgG1-Fc
 CAL04 Mock -Titration
1/2
 1/4
1/8
1/16
1/32
 1/64
 1/128
